# Defensive nymphs in the water-repellent gall of the social aphid *Colophina monstrifica* (Hemiptera: Aphididae: Eriosomatinae)

**DOI:** 10.1101/2021.05.20.443943

**Authors:** Keigo Uematsu, Shigeyuki Aoki, Man-Miao Yang

## Abstract

The aphid *Colophina monstrifica* forms woolly colonies with sterile soldiers on the secondary host *Clematis uncinata* in Taiwan. However, the gall or primary-host generation of *C. monstrifica* has not been found to date. We successfully induced galls of the species on trees of *Zelkova serrata* through attaching its eggs onto the trees, and also found a few naturally formed galls on another *Z. serrata* tree. The identity of the aphids was confirmed by examining their morphology and mitochondrial DNA sequences. First- and second-instar nymphs in the galls exhibited attacking behavior toward artificially introduced moth larvae. Observations with a scanning electron microscope revealed that the gall inner surface was densely covered with minute trichomes. This indicates the water repellency of the inner surface, and strongly suggests that young nymphs of *C. monstrifica* dispose of honeydew globules outside the gall, as known in the congener *C. clematis*.

## INTRODUCTION

Species of the aphid genus *Colophina* (Eriosomatinae; Eriosomatini) exhibit facultative host-alternation between the primary and secondary hosts. On the primary host plant, *Zelkova serrata* (Ulmaceae), a foundress (or fundatrix) forms a gall on the leaf, in which she produces offspring by parthenogenesis and young nymphs perform colony defense against intruding predators (Aoki 1980). The gall inner surface of *C. clematis* is densely covered with trichomes, which enables young nymphs to perform gall cleaning by pushing globules of honeydew out of the gall (Uematsu *et al*. 2018). This water-repellent structure can be regarded as an “extended phenotype” of the aphids in the gall (Uematsu *et al*. 2018; see also Stone & Schönrogge 2003; Kutsukake *et al*. 2019). On the secondary host plants, *Clematis* spp. (Ranunculaecae), they form dense, woolly colonies, where sterile first-instar nymphs called “soldiers” perform colony defense. Sterile soldier nymphs have been found in *C. clematis* (Aoki 1977a), *C. arma* (Aoki 1977b), *C. monstrifica* (Aoki 1983) and *C. clematicola* (Akimoto 1998). Soldiers of the first three species are characterized by their enlarged fore and mid legs, with which they cling tightly to a predator and sting it with their stylets (Ijichi *et al*. 2005).

The gall or primary-host generation of *Colophina* has been recorded in three Japanese species, *C. clematis, C. arma* and *C. clematicola* (Aoki 1980; Aoki & Kurosu 2000). However, galls of *Colophina* (*C. arma* and *C. clematicola* in particular) are rarely found under natural conditions, despite the fact that trees of their primary host, *Z. serrata*, are common in Japan (Kurosu & Aoki 1991; Aoki & Kurosu 2000). Since some young nymphs produced on the secondary host plant overwinter on buds near the ground (Aoki *et al*. 1997) or in crevices of the bark of lignified stems of *Clematis* (Aoki 1977a, 1980; Blackman & Eastop 2021), the aphids can continue their life cycles without returning to the primary host. By collecting its sexuparae from the secondary host *Clematis terniflora* and transferring them to a potted tree of *Z. serrata*, Aoki and Kurosu (2000) succeeded in inducing a few galls of *C. clematicola*, which had been unknown before.

*Colophina monstrifica* is known from mountainous areas of Taiwan (Aoki 1983). The species forms dense, woolly colonies on stems of the secondary host *Clematis uncinata* (reported as “*C. floribunda*” in Aoki (1983)), an evergreen vine, and produces winged sexuparae in autumn. *Zelkova serrata* is widely distributed in mountainous areas of Taiwan, where galls of *C. clematis* are rather commonly found (Bo-Fei Chen 2007, unpublished Master Thesis, National Chung Hsing University). However, galls of *C. monstrifica* have not been found to date. Here, we report that *C. monstrifica* also forms galls on *Z. serrata* in Taiwan. We succeeded in inducing galls of the species on trees of *Z. serrata*, and also found a few galls under natural conditions. As in *C. clematis*, the inner surface of its galls was covered by dense trichomes. In addition, we show that first- and second-instar nymphs of *C. monstrifica* perform altruistic colony defense against intruding predators.

## MATERIALS AND METHODS

### Collection of galls

Large colonies of *Colophina monstrifica* were found on *Clematis uncinata* at Huisun Experimental Forest Station (24°05’17”N, 121°02’08”E), Nantou County, Taiwan, in November 2013. We collected many winged adults (sexuparae) from the colonies, and confined approximately 100 sexuparae in each of three clear plastic containers with bundles of bark pieces of *Z. serrata*, expecting that the sexuparae would produce sexuals and that the sexual females would lay eggs between the bark pieces. (Sexual nymphs of Eriosomatinae mature into adults without taking food.) The containers were kept outdoors. On 23 December 2013, these bark bundles were attached to three trees of *Z. serrata* at the coffee plantation of Huisun Experimental Forest Station (hereafter “Coffee Plantation”). In the following spring, on 18 and 28 April 2014, we found 12 galls of *C. monstrifica* formed on leaves of two of the three *Zelkova* trees (Fig. 1). We also found seven galls of *C. monstrifica* on leaves of another tree of *Z. serrata*, to which we had not attached the bark bundles, near the lodges of Huisun Experimental Forest Station (24°05’22”N, 121°02’05”E, hereafter “Lodge”). These galls were brought to the laboratory at Chung Hsing University, Taichung, and subjected to the following behavioral experiments. Three galls were preserved in 70 or 85% ethanol together with galled leaves to investigate the gall inner structure.

**Figure 1.**
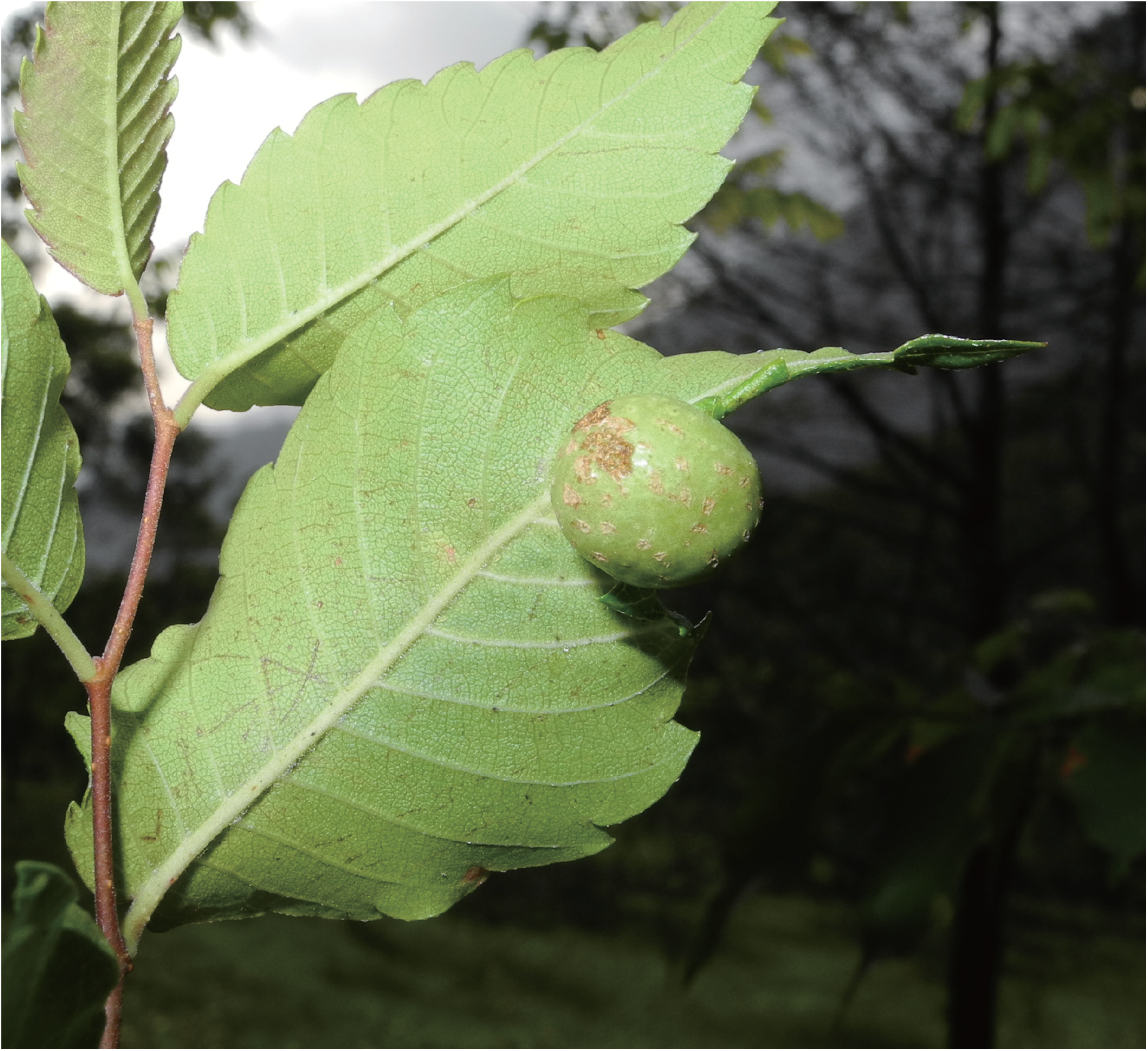
A gall of *Colophina monstrifica* formed on a leaf of *Zelkova serrata*.

### Attacking behavior against predators

To test attacking behavior of aphids inside the galls, we used young caterpillars of *Assara formosana* (Pyralidae) found in a gall of the aphid *Ceratoglyphina styracicola* as a model predator, because caterpillars of some *Assara* species are known to prey on aphids in galls (Aoki & Kurosu 2010). One caterpillar was introduced to each of four galls of *C. monstrifica*. Ten minutes later, the galls were cut open with a razor and the caterpillars were taken out of the galls, and aphids attacking the caterpillars were observed under a dissecting microscope. We also tapped a few young nymphs in a cut gall with a fine brush, and video-recorded their reaction.

### Morphological examination

Collected aphids, including those used in the experiment mentioned above, were preserved in 70 or 85% ethanol. Some of them were heated in 10% KOH solution, stained with acidic fuchsine or Evan’s blue, dehydrated through an ethanol-xylene series, and mounted on microscope slides in balsam or Mount-Quick (Daido Sangyo). The mounted individuals were photographed using a digital camera (Nikon 1) attached to a light microscope. Some slide-mounted aphids of *C. monstrifica* used in this study are deposited as voucher specimens in the collections of Department of Entomology, National Chung Hsing University, Taichung, Taiwan.

### Gall inner structure

After removing all aphids from their galls, the three gall-harboring *Z. serrata* leaves kept in ethanol were first transferred to 50% ethanol, then to FAA (formaldehyde 3.7% and acetic acid 5% in 50% ethanol), dehydrated through an ethanol series and dried. The dried samples were observed with a scanning electron microscope (SEM) and photographed. Density and length of trichomes in a 0.5 × 0.5 mm square area of the surface were measured based on the photographs using ImageJ (https://imagej.nih.gov/ij/). Statistical significance between the gall inner surface and the underside of the same leaf was analyzed using the linear mixed model (*lmer* function in the *lme4* package in R v. 3.3.3 (R Core Team 2017)) with gall identity treated as a random factor.

### Molecular phylogenetic analysis

Total DNA was extracted from three aphids of *C. monstrifica* fixed in ethanol: one collected from an artificially induced gall, another from a natural gall, and the other from a colony on *C. uncinata*. A mitochondrial DNA fragment (ca. 1.6 kb) containing small subunit rRNA, tRNA-Val, and large subunit rRNA genes was amplified by PCR, as described in Aoki *et al*. (2018), and sequenced. The DNA sequences are deposited in the DNA Data Bank of Japan (DDBJ) (accession no. LC626871). These DNA sequences and those of *C. clematis* (DDBJ/EMBL/GenBank accession no. AF275224.1) and *Eriosoma lanigerum* (accession no. NC_033352.1) were subjected to molecular phylogenetic analyses. *Ceratovacuna nekoashi* (Hormaphidinae, Cerataphidini) was used as an outgroup (accession no. AB035879.1). Multiple alignment of the nucleotide sequences was generated using MAFFT (Katoh & Standley 2013). The GTRLJ+LJG model was selected as the nucleotide substitution model using the program jModelTest2 (Darriba *et al*. 2012) based on AIC. A maximum likelihood phylogenetic tree was generated using RAxML (Stamatakis 2014). Bootstrap tests were performed with 1,000 replications.

## RESULTS

### Galls of *C. monstrifica*

Twelve globular galls (Fig. 1) were formed on the leaf of two *Z. serrata* trees, to which the bark bundles with eggs of *C. monstrifica* had been tied in the previous December. No galls were found on other *Z. serrata* trees, to which the bark bundles had not been tied, in Coffee Plantation.

A few galls were also found on a tree of *Z. serrata* in Lodge, near the collection site of the free-living colonies on *C. uncinata*. There was no difference in shape between the experimentally induced galls and the natural galls. The long distance from the tree to Coffee Plantation (about 2.2 km away in a straight line) and its close proximity to the colonies on *C. uncinata* (about 180 m away) suggest that these galls were formed naturally, possibly by grandoffspring of the sexuparae which had flown from colonies on the nearby secondary host plants.

### Colony composition

Table 1 shows the composition of inhabitants for nine galls of *C. monstrifica*. The collected galls contained a high proportion of first- and second-instar young nymphs, and a small number of nymphs with wing buds, but no winged adults. All foundresses survived and still contained embryos inside.

### Attacking behavior

When tapped with a fine brush, young nymphs in the gall exhibited an aggressive response by clutching the brush using their forelegs (Movie S1). To test their attacking response to predators, a caterpillar of *A. formosana* was introduced into four galls (Galls #5, 6, 7, 9 in Table 1). One to five (average = 2.25) first-instar nymphs and one to six (average = 2.7) second-instar nymphs clung onto the caterpillar and stung it with their stylets (Fig. 2). We confirmed under a dissecting microscope that their stylets were inserted in the body of the caterpillar, and their claws penetrated the skin. Thirty minutes later, all four caterpillars were completely immobilized.

**Figure 2.**
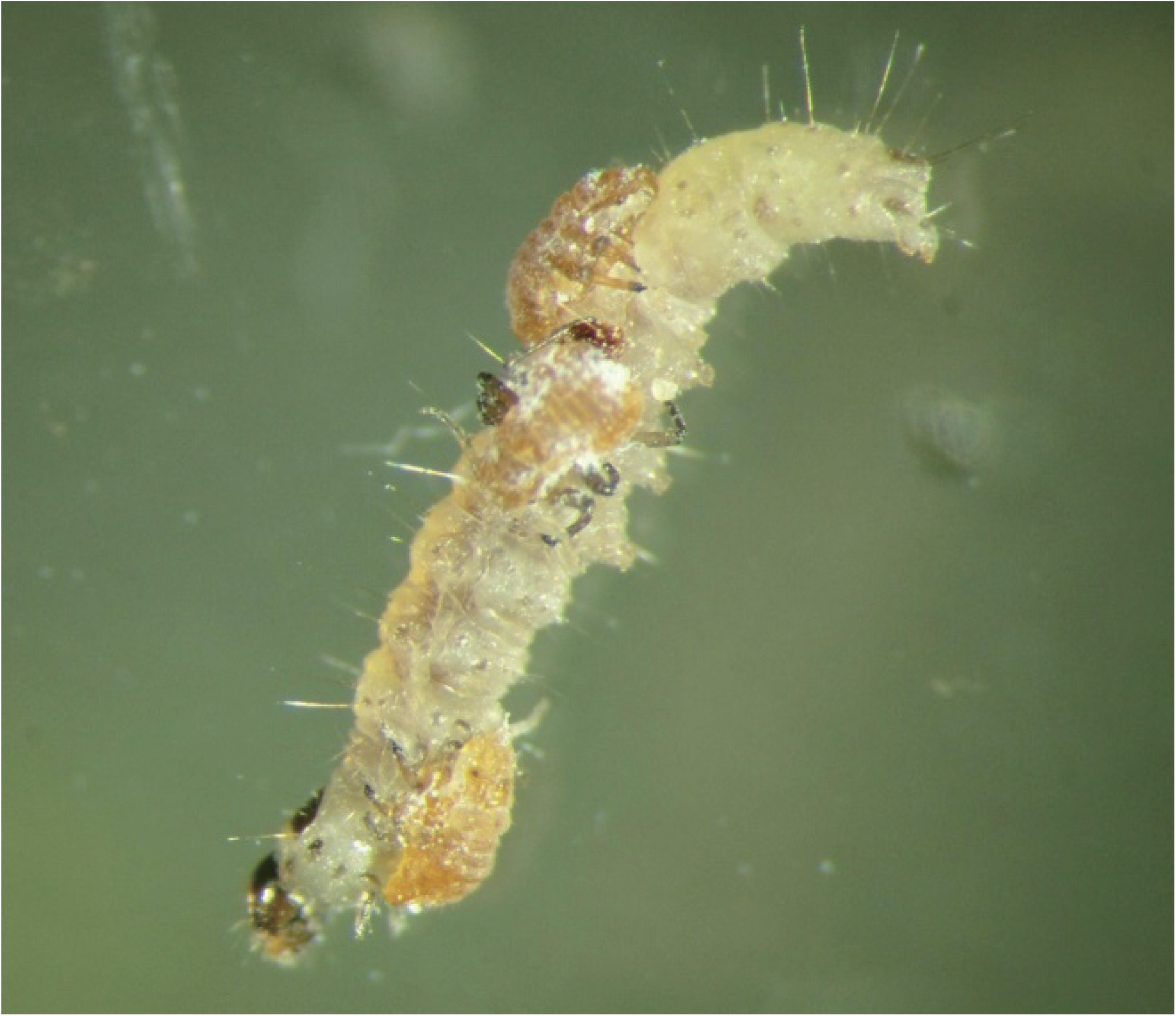
Young nymphs of *Colophina monstrifica* attacking a larva of the moth *Assara formosana* introduced into their gall.

### Morphology of gall inhabitants

Among the gall inhabitants of *C. monstrifica*, the first- and second-instar nymphs (Fig. 3) had well-developed fore and mid legs with large, strongly curved claws. In particular, fore and mid legs of the first-instar nymphs were distinctly thickened (Fig. 3a). As in *C. clematis* and *C. arma* (Aoki 1980), the first-instar nymphs (Fig. 3a) were discriminated from the second-instar nymphs (Fig. 3b) by the long, usually capitate dorsoapical setae on the second segment of each tarsus, and by the lack of short, spine-like seta (sense peg) on the first tarsal segment. No remarkable morphological differences were observed between attacking and non-attacking individuals, indicating that the first- and second-instar nymphs are monomorphic defenders.

**Figure 3.**
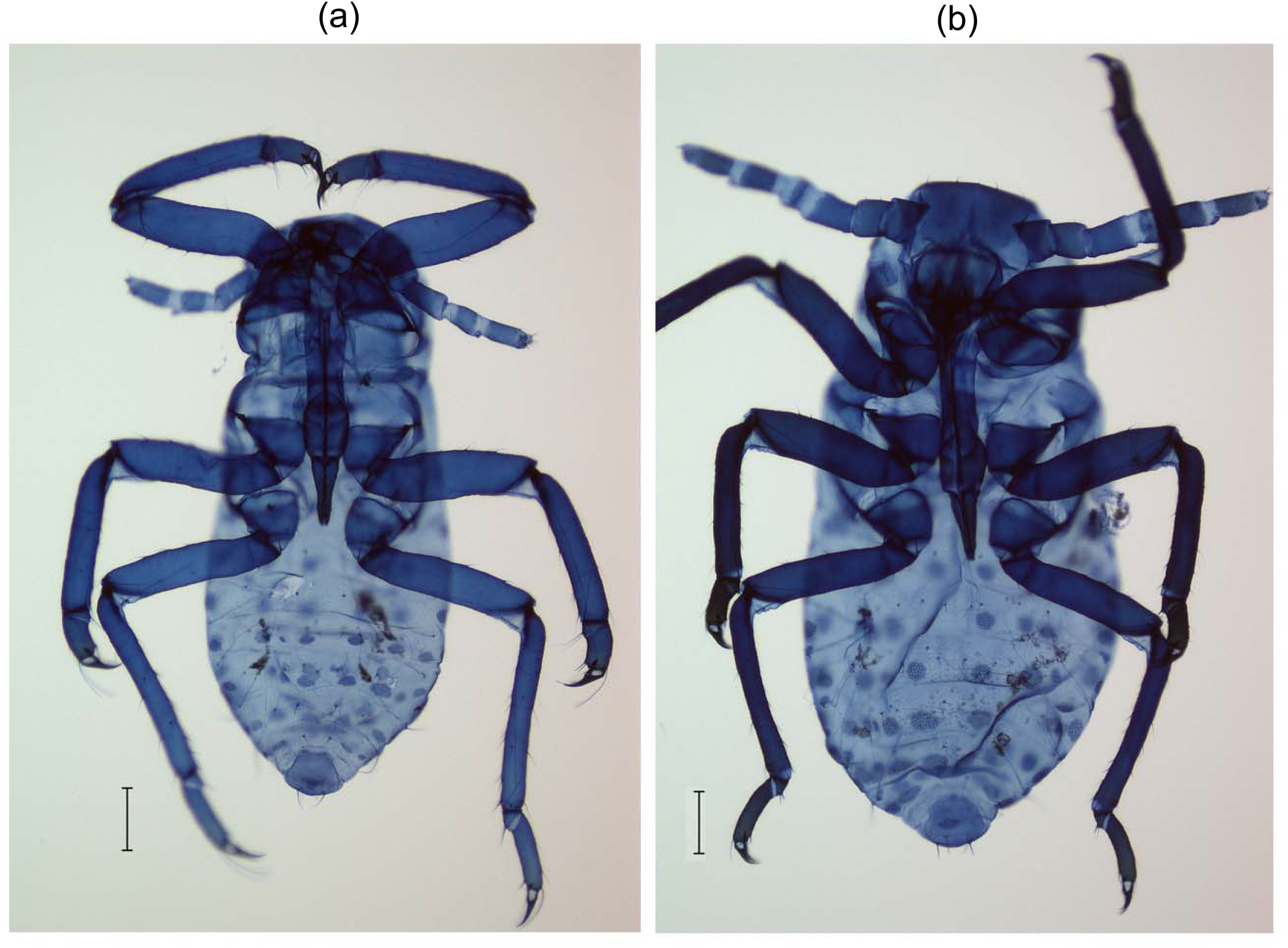
Gall generation of *Colophina monstrifica*: (a) first-instar nymph (from a gall collected on 18 April 2014); (b) second-instar nymph (from a gall collected on 28 April 2014). Scale bars represent 100 μm.

These gall inhabitants were morphologically distinguishable from the primary host generation of *C. clematis*, which has been the only known *Colophina* species that forms a globular gall on the leaf of *Z. serrata* in Taiwan. The first-instar nymphs of *C. monstrifica* had a pair of small, half ring-like cornicles on the flat tergite (Fig. 4c), whereas the first instar nymphs of *C. clematis* have distinctly protruded cornicles (Fig. 4d). In addition, the apex of each antenna was rounded in the first instar nymphs of *C. monstrifica* (Fig. 4a), while the apex is conical in those of *C. clematis* (Fig. 4b).

**Figure 4.**
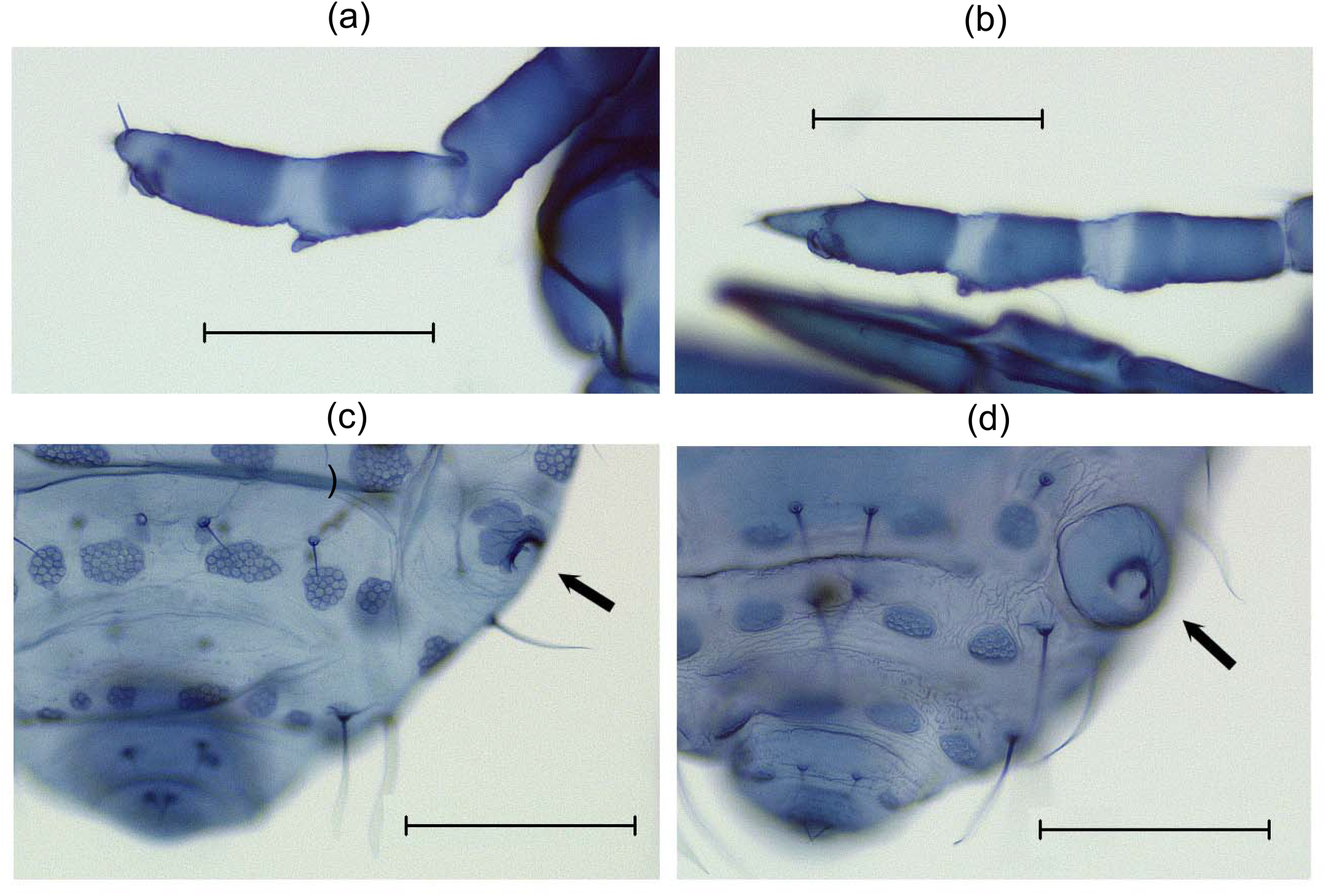
First-instar nymphs of *Colophina monstrifica* and *C. clematis* produced in the galls on *Zelkova serrata*: (a) apical antennal segments of *C. monstrifica*; (b) apical antennal segments of *C. clematis*; (c) posterior abdominal tergites of *C. monstrifica*; (d) posterior abdominal tergites of *C. clematis*. The right cornicle is indicated by an arrow in (c) and (d). The photographed nymphs (a, c) of *C. monstrifica* were collected on 18 April 2014, and the photographed nymph (b, d) of *C. clematis* was collected in Miaoli, Taiwan, on 15 May 2002. Scale bars represent 100 μm.

### Molecular phylogenetic analysis

The DNA sequences (1,579 bp) of the three individuals of *C. monstrifica* (collected from an artificially induced gall, from a natural gall, and from a colony on *C. uncinata*) were completely identical with each other. The molecular phylogenetic analysis including other eriosomatine aphids also showed that the three samples were clearly distinct from *C. clematis* (Fig. 5), confirming our identification of *C. monstrifica*.

**Figure 5.**
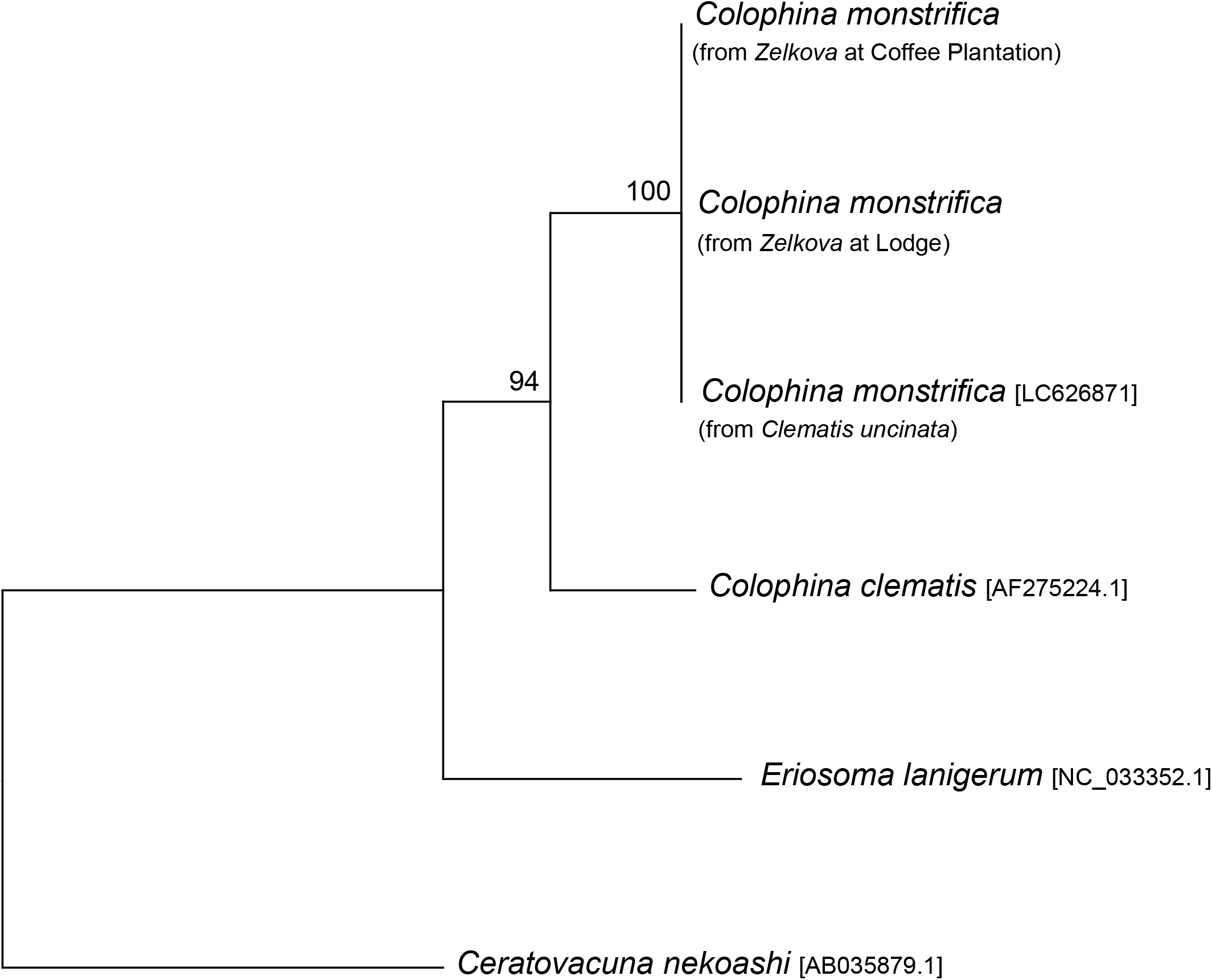
Molecular phylogenetic analysis of eriosomatine aphid species. The maximum likelihood phylogeny inferred from aligned 1,572 nucleotide sites of the mitochondrial rRNA gene is shown. Bootstrap probability in percentage is shown at the nodes. The DDBJ/EMBL/GenBank accession number for each DNA sequence is indicated in square brackets.

### Trichomes on the inner surface of galls

The observations using an SEM revealed that the inner surface of the galls of *C. monstrifica* was densely covered with tiny trichomes (Fig. 6). The trichome density was 298.3 ± 44.7 / mm^2^ (n = 12), which was significantly higher than the trichome density on the underside of the same leaf (16.7 ± 3.7 / mm^2^) (n = 12, χ^2^ = 74.7, df = 1, *P* < 0.001). On the other hand, trichomes on the inner surface of the galls were 82.2 ± 30.0 μm (n = 30) in length and shorter than those on the underside of the leaf, which were 105.0 ± 45.3 μm (n = 30, χ^2^ = 4.97, df = 1, *P* = 0.026). The high trichome density on the gall inner surface was comparable to that of *C. clematis* (221.7 trichomes / mm^2^ on average) found in a previous study (Uematsu *et al*. 2018).

**Figure 6.**
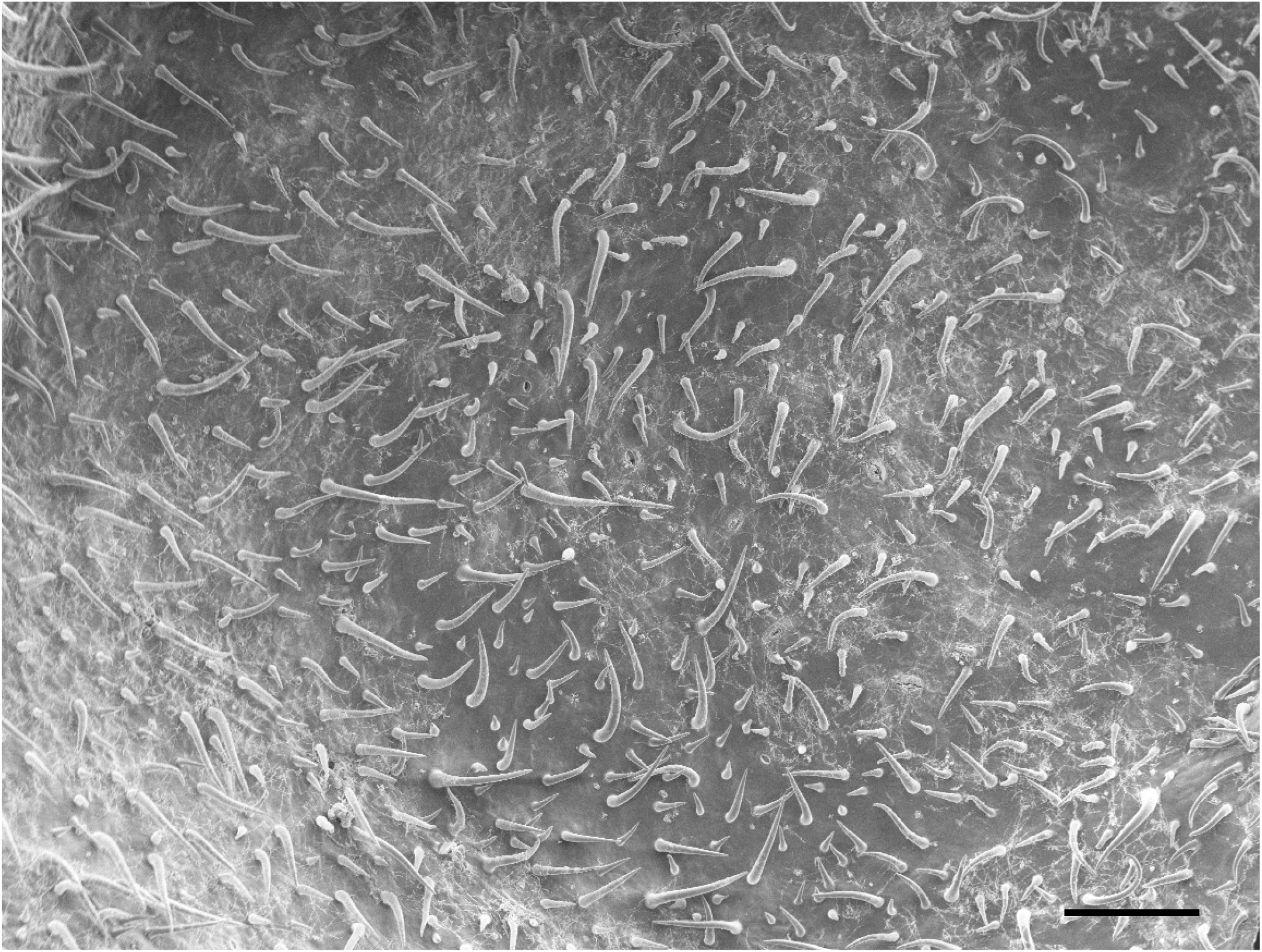
Trichomes on the inner surface of a gall of *Colophina monstrifica*. Scale bar represents 200 μm.

## DISCUSSION

We collected naturally occurring and artificially induced galls of *C. monstrifica* on *Zelkova serrata* in Taiwan, and confirmed the identity of aphids in the galls by subsequent morphological and molecular analyses. The aphids were distinct in morphology from *C. clematis*, the only congener known in Taiwan. The molecular analyses also revealed the identity of the mitochondrial DNA sequences between the aphids in these galls on *Z. serrata* and those collected from the secondary host *Clematis uncinata*. Although migration of winged adults from the gall to *C. uncinata* has not been observed, these results indicate that, like other *Colophina* species, *C. monstrifica* has a life cycle with host alternation between *Zelkova* and *Clematis*. We found colonies of *C. monstrifica* on *C. uncinata* at Huisun Experimental Forest Station also on 28 March 2011. This indicates that the host alternation of *C. monstrifica* is facultative, or that its colonies can persist on the clematis throughout the year.

We confirmed that first- and second-instar nymphs of *C. monstrifica* attack predators introduced in their gall. Their attack using the claws and stylets can immobilize and kill such potential predators as pyralid larvae. In addition, we showed that the inner surface of galls of *C. monstrifica* was densely covered with minute trichomes. The galls of *C. clematis* and *C. arma* also have dense trichomes on the inner surface. It is known that these trichomes collect wax particles produced by aphids, make the inner surface water repellent and facilitate the formation of small globules of honeydew covered with the wax. Young nymphs of *C. clematis* actively dispose of honeydew globules outside their gall through a small opening (Uematsu *et al*. 2018). Although we did not directly observe honeydew-disposing behavior in this species, the microstructure of the inner surface of the gall strongly suggests that young nymphs of *C. monstrifica* also dispose of honeydew globules.

There are many species of Eriosomatinae and Hormaphidinae whose gall or primary-host generations are unknown (Blackman & Eastop 2021). It is not clear whether these species lost their primary-host generations irreversibly or still retain the ability to induce galls on the primary host. In this study, we obtained eggs of *C. monstrifica* by collecting sexuparae from the secondary host, attached them to its presumed primary host, and successfully induced galls of the species, which had been unknown before. This method may be applied to other species for elucidating their life cycles.

## ACKNOWLEDGMENTS

We thank Utako Kurosu for her advice on how to collect eggs of *Colophina* spp., and Wan-Zon Yang for offering us a sample of *Colophina clematis*. We also thank Chun-I Chiu, Yi-Chuan Lee, and Sheng-Feng Lin, and Chang-Ti Tang for their help during the survey. K. U. was supported by the Sumitomo Foundation (130972) and JSPS Overseas Research Fellowships (24-596). The collection of aphids at Huisun Experimental Forest Station was permitted by the Experimental Forest Management Office, NCHU (nos. 1020000228 and 1030000107).

## SUPPORTING INFORMATION

Additional Supporting Information may be found online in the Supporting Information section at the end of the article.

**Movie S1**. Attacking behavior of young nymphs of *Colophina monstrifica* toward a fine brush in a cut gall.

